# Brain functional connectivity correlates of response in the 7.5% CO_2_ inhalational model of generalized anxiety disorder: a pilot study

**DOI:** 10.1101/823617

**Authors:** Nathan T.M. Huneke, M. John Broulidakis, Angela Darekar, David S. Baldwin, Matthew Garner

**Author notes:** Contributed equally (David S. Baldwin, Matthew Garner). Corresponding author: Nathan TM Huneke, University Department of Psychiatry, Academic Centre, College Keep, 4-12 Terminus Terrace, Southampton, SO14 3DT, UK.

## Abstract

The 7.5% CO_2_ inhalational model (‘CO_2_ challenge’) can be used to explore potential treatments for generalized anxiety disorder. However, it remains unknown how inter-individual variability in the functional architecture of negative affective valence systems might relate to the anxiogenic response to CO_2_ challenge. In this pilot study, we explored how connectivity in systems associated with processing potential threat (“anxiety”) correlated with behavioural measures of anxiety following prolonged CO_2_ inhalation.

The negative affective valence system was identified using a passive emotional face perception task. Spherical regions of interest were created from peak voxels of significant brain activation when 100 young adult participants viewed emotional faces compared with black and white concentric circles during a functional MRI scan. Using these regions of interest, generalized psychophysiological interaction (gPPI) analysis was undertaken to explore task-evoked functional connectivity in a separate group of 13 healthy volunteers. Within 7 days of the scan, these participants underwent CO_2_ challenge and results from the gPPI analysis were correlated with CO_2_ outcome measures.

Exposure to CO_2_ challenge significantly increased subjective anxiety, negative affect, systolic blood pressure and heart rate. Functional connectivity between the ventromedial prefrontal cortex and right amygdala was positively correlated with heart rate. Increased connectivity between the vmPFC and the right amygdala, and decreased connectivity between the midcingulate cortex (MCC) and the left amygdala, correlated with subjective anxiety during CO_2_ challenge.

Response to CO_2_ challenge was related to task-evoked functional connectivity between regions known to be important in processing potential threat. Further studies are required to assess whether this translates into clinical populations. Measures of functional connectivity within emotional processing networks could be potential biomarkers to enable stratification of healthy volunteers, and to examine correlates of response, in trials using experimental medicine models.

## Introduction

Experimental medicine models in healthy volunteers can be a cost-effective and timely approach to explore potential novel treatments for psychiatric disorders. An example is the 7.5% CO_2_ inhalational model of generalized anxiety disorder (GAD), in which healthy volunteers inhale air “enriched” with 7.5% CO_2_ (‘CO_2_ challenge’). This model mimics the subjective, autonomic and neuropsychological features of GAD (Bailey et al., 2005; Garner et al., 2011; Garner et al., 2012). Anxiety induced in this model is responsive to standard pharmacological and psychological treatment (Ainsworth et al., 2015; Bailey et al., 2007; Pinkney et al., 2014). Therefore, this model provides an approach for testing potential treatments for GAD at a ‘proof of concept’ stage, prior to embarking on time-consuming and costly phase II/III clinical trials (Bailey et al., 2011a; Bailey et al., 2011b; Poma et al., 2014; Baldwin et al., 2017).

However, there is variation in how healthy volunteers respond to CO_2_ challenge. Volunteers scoring highly in trait anxiety and anxiety sensitivity measures experience increased subjective and physiological responses to CO_2_ challenge, when compared with those with lower scores (Fluharty et al., 2016; Olatunji et al., 2009). Further, both increased anxiety sensitivity and CO_2_ challenge reactivity predict subsequent development of clinically typical anxiety (Schmidt and Zvolensky, 2007), though how these factors potentially interact to predict the development of pathology remains unclear. Both increased anxiety sensitivity and 7.5% CO_2_ inhalation increase selective attention biases to threatening stimuli (Garner et al., 2011; Lees et al., 2005; Hunt et al., 2006; Keogh et al., 2001). This suggests that the function of ‘negative affective valence systems’ (defined by NIMH research domain criteria https://www.nimh.nih.gov/research/research-funded-by-nimh/rdoc/index.shtml), particularly in the constructs of acute and potential threat, might be important.

A key anatomical node identified in the negative affective valence system is the amygdala. Amygdala hyperactivity during the processing of negative emotion is a commonly reported finding in patients with GAD (Etkin and Wager, 2007; Mochcovitch et al., 2014), and exhibits abnormal functional connectivity with regions important in emotional processing at rest and during the perception of fearful faces (Etkin et al., 2009; Prater et al., 2013; Rabany et al., 2017). Similarly, functional connectivity of the amygdala also relates to measures of anxiety in healthy volunteers. For example, resting state functional connectivity between the amygdala and dorsomedial prefrontal cortex (PFC) is positively correlated with anxiety, and amygdala/ventromedial PFC connectivity negatively correlated (Kim et al., 2011). Furthermore, trait anxiety was negatively correlated with connectivity between the midcingulate cortex (MCC) and the left amygdala when viewing negative pictures (Kienast et al., 2008). It is therefore possible that inter-individual differences in the functional architecture of negative valence systems might be a biomarker of prospective subjective and physiological response to CO_2_ challenge.

We carried out a pilot study to explore how functional connectivity within negative affective valence networks is associated with response to CO_2_ challenge in healthy volunteers. To probe functional connectivity of these networks under different conditions, we used a validated passive emotional face perception task known to activate the amygdala (Schneider et al., 2011; Grosbras and Paus, 2006). Through generalized psychophysiological interaction analysis, we identified task-evoked changes in functional connectivity and correlated these findings with CO_2_ outcome measures. Our aim was to identify whether there were any signals of interest to inform further hypotheses and future studies.

## Method

### Ethics statement

This study was reviewed and approved by the Ethics and Research Governance Office at the University of Southampton (reference: 27440) All participants provided written, informed consent prior to taking part.

### Participants

We recruited 13 healthy volunteers (mean age 23.38 ± 4.27 years, 8 females) through adverts placed on the University campus and in the local community. Participants who responded were invited to attend a screening interview to determine eligibility. We excluded participants if they had current or lifetime history of psychiatric illness (as assessed by the Mini International Neuropsychiatric Interview for DSM-IV - MINI (Sheehan et al., 1998)), chronic physical illness, or alcohol or drug dependence. We also excluded participants if they had used any medication, alcohol or illicit drugs in the previous 8 weeks, if they were regular smokers, if there was any contraindication to MRI scanning, or if body mass index was less than 18 or greater than 28 kg/m^2^.

### Study Design

#### Trait anxiety measures

Following the screening interview, eligible participants completed the following baseline measures of trait anxiety: the State-Trait Anxiety Inventory (STAI-trait) (Spielberger et al., 1983), the Anxiety Sensitivity Index (ASI) (Peterson and Reiss, 1992), and a modified version of the Generalized Anxiety Disorder Screener (GAD-7) (Spitzer et al., 2006), where each question was represented by a visual analogue scale ranging from “Not at all” to “Nearly every day”.

#### MRI scanning session

Participants attended University Hospital Southampton, Southampton, UK, for a functional magnetic resonance imaging (fMRI) scan. During the scan, we asked participants to complete a validated passive emotional face perception task (Grosbras and Paus, 2006). In brief, participants passively viewed short black and white video clips of actors making emotional facial expressions. The actors’ faces always started from a neutral expression, and then either turned angry, or happy, or the actor made a neutral movement with no emotional content. These stimuli were arranged into 18-second blocks of faces portraying the same emotion. These blocks were interspersed with an 18-second control condition of expanding and contracting black and white concentric circles. Altogether, participants viewed 4 blocks of happy faces, 4 blocks of angry faces, 4 blocks of neutral faces, and 12 blocks of the control condition (total 24 blocks). We chose this task as it has been shown to reliably activate the amygdala (Schneider et al., 2011), and would therefore be a good probe of negative affective valence network function.

#### CO_2_ challenge

Within 7 days of the MRI scan, participants completed two 20-minute inhalations of normal air and subsequently air enriched with 7.5% CO_2_ (21% O_2_, balance N_2_) administered through an oronasal face mask, under single-blind conditions. We measured heart rate, blood pressure (with an automated sphygmomanometer: Omron-M6, Medisave, UK) and subjective mood and anxiety at pre-test baseline and immediately following each 20-minute inhalation of air and 7.5% CO_2_. We measured subjective mood with the Positive and Negative Affect Schedule (PANAS) (Watson, 1988). Subjective state anxiety was measured with a modified version of the GAD-7, where each question was represented by a visual analogue scale ranging from “Not at all” to “All of the time”. For both the PANAS and the GAD-7, we asked participants to answer based on the peak anxiety they experienced over the previous 20 minutes.

### Image Acquisition

Images were acquired using a 20 channel head coil, on a 3T Siemens Skyra MRI scanner (Siemens Healthineers Limited), at University Hospital Southampton, Southampton, UK. T1-weighted (MP-RAGE) anatomical images were acquired for registration purposes (voxel size 1 × 1 × 1mm; TR 2200ms; TE 2.45ms; flip angle 8 degrees; 176 sagittal slices). Functional MR images were obtained with a T2*-weighted single-shot gradient echo, echo planar imaging sequence (voxel size 2.5 × 2.5 × 2.5mm; TR 2500ms; TE 30.0ms; parallel imaging method: GRAPPA with acceleration factor 4; flip angle 90 degrees; FOV 235mm × 235mm; 45 slices in an oblique orientation, and an interleaved acquisition at 2.5mm slice thickness with no gap). The single scanning session consisted of 178 volumes synchronised with the onset of the experimental task.

### Image Pre-processing

We used FEAT (FMRI Expert Analysis Tool) version 5.0.8, part of FSL (FMRIB’s Software Library, www.fmrib.ox.ac.uk/fsl) for image preprocessing. Preprocessing steps included slice time correction, motion correction using MCFLIRT (Jenkinson et al., 2002), brain extraction, spatial smoothing with a gaussian filter with a full width at half maximum kernel of 5mm, and application of a high pass filter with a 160s cutoff. None of our participants had movement greater than 2mm translation or 2 degrees rotation. We carried out linear registration of functional images to high resolution structural images (T1 MP-RAGE) and then to standard space images (T1-weighted Montreal Neurological Institute template) with 12 degrees of freedom, using FLIRT (Jenkinson et al., 2002; Jenkinson and Smith, 2001). Registration from high resolution structural to standard space was then further refined with FNIRT nonlinear registration (Andersson et al., 2007a; Andersson et al., 2007b).

### Statistical analyses

#### Analysis of CO_2_ challenge

We undertook statistical analyses of CO_2_ challenge outcomes using the SPSS software package (IBM SPSS Statistics for Windows version 24.0; IBM Corp., Armonk, NY). We assessed response through ANOVA with repeated measures for each outcome measure with time as the within-subject factor (pre-test baseline, post-air and post-CO_2_). We used Greenhouse-Geisser correction where sphericity assumptions were violated.

#### Functional MRI analyses

We first performed whole brain analysis of the contrast All Faces > Control to check whether participants had responded to the task as expected. At the individual level, we applied a general linear model to the fMRI timeseries for each task condition convolved with a single-gamma canonical haemodynamic response function. Higher level analysis was carried out using a mixed effects statistical model (FLAME stage 1 and stage 2 (Beckmann et al., 2003; Woolrich, 2008; Woolrich et al., 2004)). We applied voxel-wise control of family-wise error with a (corrected) significance threshold of *p* = 0.05.

We explored task-modulated functional connectivity through generalized psychophysiological interaction (gPPI) analysis using the CONN toolbox (www.nitrc.org/projects/conn, RRID:SCR_009550). In brief, this analysis involves calculating a separate multiple regression model for target regions of interest, each including psychological predictors (all task effects convolved with a canonical haemodynamic response function) and physiological predictors (average BOLD timeseries from seed region), and the interaction of these (their product). We carried out generalized PPI as this has been shown to better capture task effects compared with standard PPI methods (McLaren et al., 2012). Due to our small sample size we restricted the analysis to 10 regions of interest (ROI) to improve signal to noise ratio. We defined these from a sample of 100 young adults who were scanned while undergoing the same passive emotional face perception task for another study carried out in the department. Spheres with radius of 6mm were created from peak coordinates in significant clusters seen in the contrast All Faces > Control in this sample (Z > 2.3, corrected *p* < 0.05) (see Table 1).

**Table 1:** Regions of interest used in the generalized psychophysiological interaction analysis

| **Anatomical Region** | **MNI coordinates** |  |  |
| --- | --- | --- | --- |
|  | **x** | **y** | **z** |
| *Right* |  |  |  |
| Amygdala | 20 | -4 | -14 |
| SMA | 4 | 10 | 60 |
| Angular gyrus | 34 | -56 | 40 |
| Thalamus | 12 | -14 | 8 |
| vmPFC | 4 | 54 | -14 |
| *Left* |  |  |  |
| Amygdala | -20 | -4 | -14 |
| Caudate | -14 | -6 | 18 |
| Superior temporal gyrus | -50 | 4 | -18 |
| MCC | -6 | 4 | 28 |
| Supramarginal gyrus | -50 | -48 | 10 |

Abbreviations: MNI, Montreal Neurological Institute; SMA, supplementary motor area; vmPFC, ventromedial prefrontal cortex; MCC, midcingulate cortex.

For each seed ROI, bivariate regression matrices were calculated, yielding standardised regression coefficients at the group level. We included motion parameters as a confound regressor at the lower level. CO_2_ challenge outcome measures were included as higher-level covariates of interest to assess the relationship between task-evoked functional connectivity and response to 7.5% CO_2_ inhalation. We report results at a threshold of *p* < 0.05 (2-tailed) with false discovery rate (FDR) correction at the level of the entire analysis (i.e. controlling for each ROI to target pair simultaneously). As this was a pilot study, we also performed exploratory analyses with an uncorrected significance threshold of *p* < 0.01 (2-tailed).

## Results

### Trait anxiety measures

At baseline, participants’ mean (±SD) trait anxiety measures were as follows: STAI-trait, 31.31 (±9.79); ASI, 7.77 (±4.36); GAD-7, 1.44 (±1.07). These results suggest our sample showed low trait anxiety and low anxiety sensitivity.

### CO_2_ challenge results

The CO_2_ challenge significantly increased subjective anxiety (F_(1,15)_ = 12.89, *p* = 0.002), negative affect (F_(1,13)_ = 7.45, *p* = 0.015), systolic blood pressure (F_(2,22)_ = 6.57, *p* = 0.007), and heart rate (F_(2,19)_ = 12.26, *p* = 0.001). There was an effect of time on positive affect (F_(1,20)_ = 19.56, *p* < 0.001), which was driven by a decrease in positive affect after the air inhalation. These results are summarised in Table 2.

**Table 2:** Summary of results of repeating measures ANOVAs analysing CO_2_ challenge results

| Outcome measure | Pre Air | Post Air | Post CO<sub>2</sub> | ANOVA |
| --- | --- | --- | --- | --- |
| GAD-7 | 1.43 ± 0.42 | 1.17 ± 0.32 | 5.14 ± 1.16 | $F_{(1,15)} = 12.89, p = 0.002,$<br>partial $\eta^2 = 0.518$ |
| PANAS Positive Affect | 25.00 ± 1.84 | 16.62 ± 0.97 | 16.62 ± 0.90 | $F_{(1,17)} = 19.56, p < 0.001,$<br>partial $\eta^2 = 0.620$ |
| PANAS Negative Affect | 12.08 ± 0.67 | 10.92 ± 0.31 | 18.53 ± 2.61 | $F_{(1,13)} = 7.45, p = 0.015,$<br>partial $\eta^2 = 0.383$ |
| Heart rate (bpm) | 74.38 ± 3.03 | 78.08 ± 3.37 | 92.15 ± 4.95 | $F_{(2,19)} = 12.26, p = 0.001,$<br>partial $\eta^2 = 0.505$ |
| Systolic blood pressure (mmHg) | 122.54 ± 4.72 | 115.23 ± 3.27 | 129.31 ± 6.62 | $F_{(2,22)} = 6.57, p = 0.007,$<br>partial $\eta^2 = 0.354$ |
| Diastolic blood pressure (mmHg) | 72.46 ± 2.46 | 70.38 ± 1.92 | 71.31 ± 2.81 | n.s. |

Values are reported as mean ± standard error. Abbreviations: GAD-7, Generalized Anxiety Disorder Screener; PANAS, Positive and Negative Affect Schedule; BPM, beats per minute.

### fMRI Analysis

#### Whole-brain analysis

We initially carried out a whole-brain analysis of task-related activation for the contrast All faces > control to ensure that participants responded to the task as expected. These results are summarised in Table 3 and Figure 1. As expected, we found significant activation in areas including right middle frontal gyrus, left orbitofrontal cortex, bilateral amygdala, and bilateral fusiform cortex (*Z* > 5.91, *corrected p* < 0.0005).

**Table 3:**
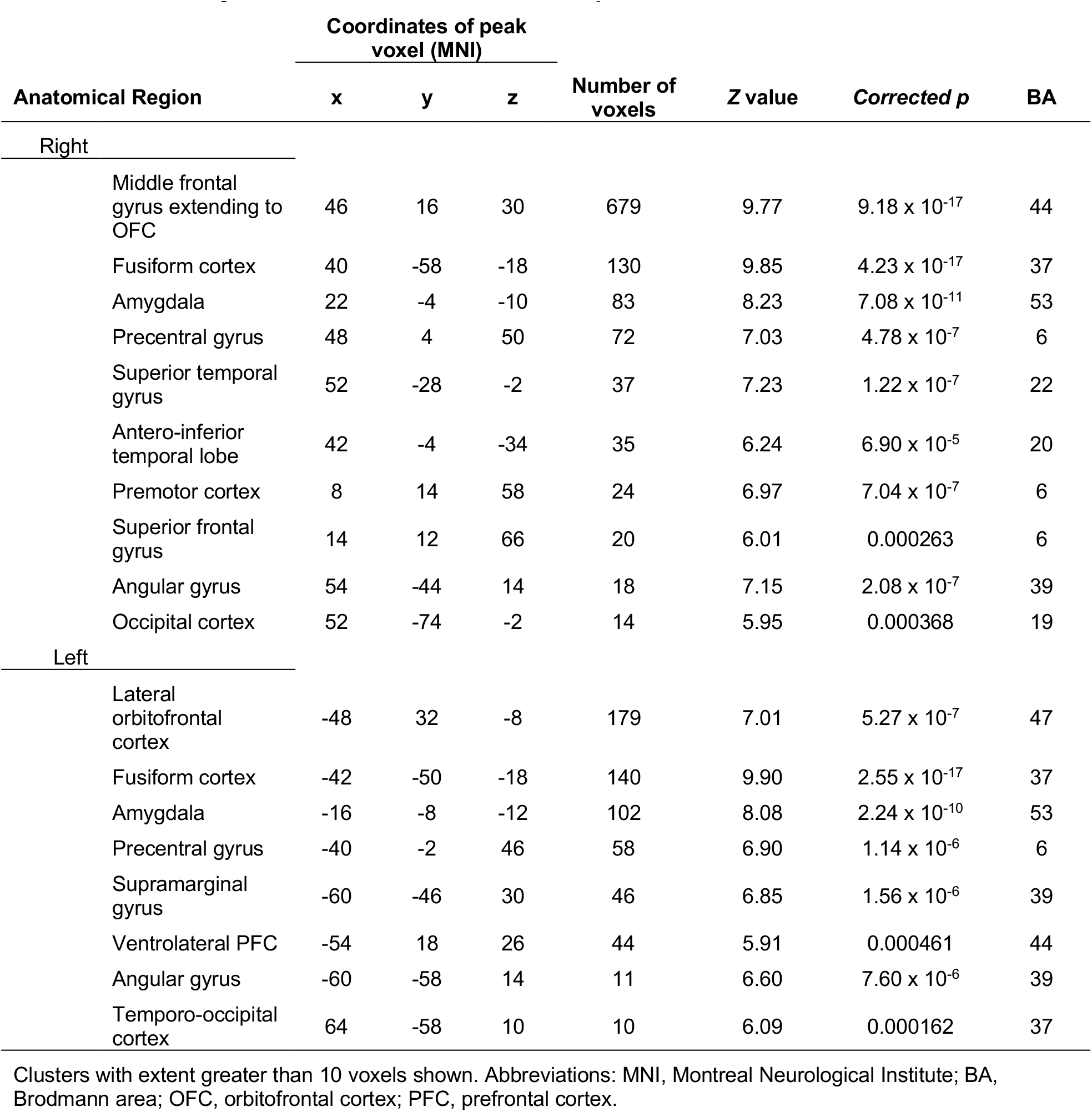
Clusters of significant activation in the passive emotional face perception task, All faces > Control. Only clusters with voxel extent >10 reported.

**Figure 1.**
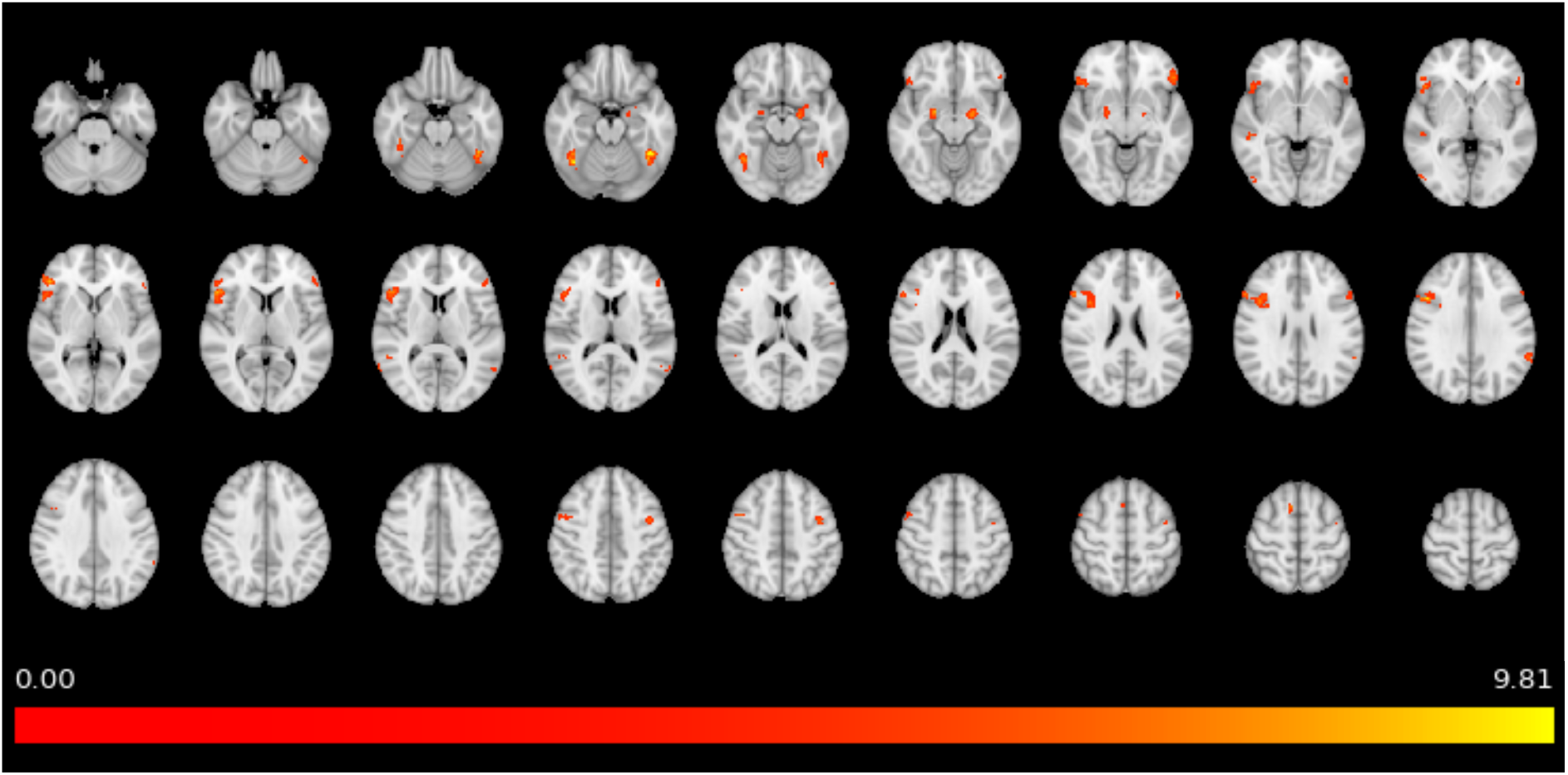
Significant activation seen on whole-brain analysis of the contrast All faces > Control in the passive emotional face perception task; *Z* > 5.91, voxelwise corrected *p* < 0.0005. Colourbar represents Z value.

#### Generalized psychophysiological interaction analysis

We then conducted generalized psychophysiological interaction analysis with 10 regions of interest. These results are summarised in Figure 2. Heart rate following CO_2_ challenge was positively correlated with functional connectivity between the ventromedial prefrontal cortex (vmPFC) and right amygdala in the contrast angry > happy (R^2^ = 0.691, t_(11)_ = 4.96, FDR-corrected *p* = 0.0039). There were no other significant relationships between baseline trait anxiety measures, CO_2_ outcome measures and task-evoked functional connectivity when correcting for multiple comparisons.

**Figure 2.**
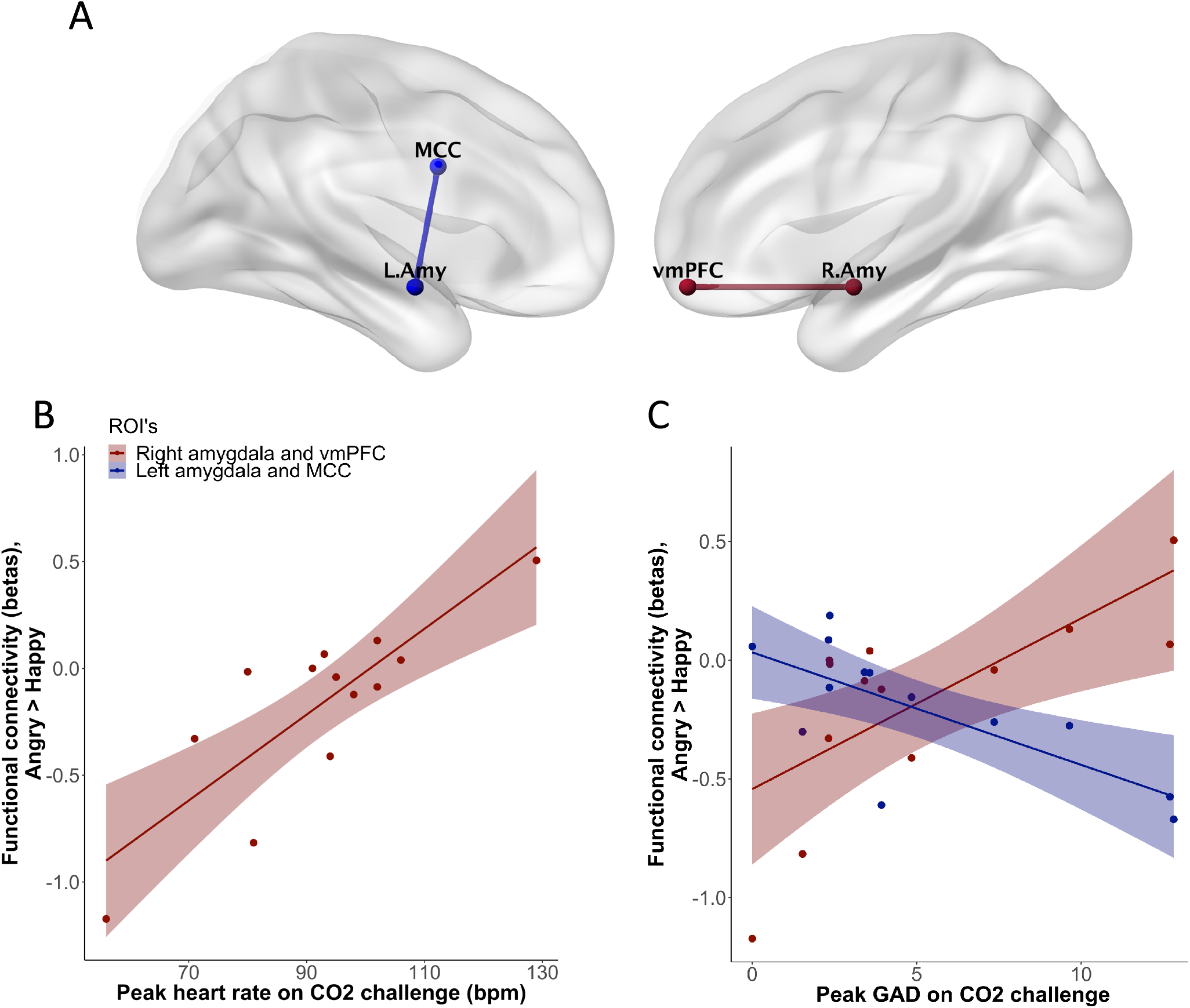
A) Task-evoked functional connectivity (contrast Angry > Happy) between right amygdala and vmPFC and between left amygdala and MCC was correlated with CO_2_ outcome measures. These connectivity results were visualized with the BrainNet Viewer (http://www.nitrc.org/projects/bnv/) (Xia et al., 2013). (B) and (C) are scatter plots showing the association of functional connectivity in these regions of interest with heart rate and peak subjective anxiety during CO_2_ challenge. Abbreviations: vmPFC, ventromedial prefrontal cortex; MCC, midcingulate cortex; bpm, beats per minute; GAD, generalized anxiety disorder screener.

We also carried out exploratory analyses with a more liberal threshold of uncorrected *p* < 0.01 to assess for trend-level results. With this threshold we found additional results, again in the contrast angry > happy. GAD-7 scores were positively correlated with functional connectivity between the vmPFC and right amygdala (R^2^ = 0.489, t_(11)_ = 3.25, *uncorrected p* = 0.0078), and negatively correlated with connectivity between the MCC and left amygdala (R^2^ = 0.527, t_(11)_ = −3.50, *uncorrected p* = 0.0050). Relationships between baseline trait anxiety measures, PANAS score, blood pressure and functional connectivity did not reach trend-level significance.

## Discussion

We performed a pilot study to explore how inter-individual variability in the functional architecture of negative affective valence networks is associated with response to the CO_2_ challenge. Consistent with previous literature, inhalation of 7.5% CO_2_ significantly increased subjective and autonomic measures of anxiety (Ainsworth et al., 2015; Bailey et al., 2005; Bailey et al., 2007; Garner et al., 2011; Garner et al., 2012). In addition, we found a significant relationship between anxiety during CO_2_ challenge and functional connectivity when viewing angry compared with happy faces. Increased connectivity between the vmPFC and the amygdala was positively correlated with increased heart rate and with subjective anxiety, while subjective anxiety was negatively correlated with connectivity between the MCC and the amygdala.

In this study, we adopted a novel approach to assess whether task-evoked functional connectivity could be a marker of anxiogenic response in an experimental medicine model of GAD. We used a task that reliably activates the amygdala to probe the functional neuroanatomy of negative affective valence systems and then correlated our findings with behavioural response in the 7.5% CO_2_ model of anxiety. We focused on a task that activates the amygdala as this region is known to be important in the generation of negative emotional experiences (Murphy et al., 2003; Phan et al., 2002). Whole-brain analysis of the overall effect of the task (All faces > control contrast) showed that regions known to be important in processing emotional faces, such as the amygdala, fusiform gyrus and the superior temporal gyrus, were activated (Schneider et al., 2011; Grosbras and Paus, 2006). This demonstrates that our participants responded to the task as expected.

Increased vmPFC-right amygdala connectivity when viewing angry compared with happy faces was significantly correlated with heart rate, and at a trend level with subjective anxiety, during CO_2_ challenge. Previous studies have shown a relationship between vmPFC activity, sympathetic tone and anxiety. For example, regional cerebral blood flow in the vmPFC during passive anticipation of shock was positively correlated with both heart rate and subjective anxiety (Simpson et al., 2001). Further, patients with vmPFC damage exhibit reduced skin conductance responses to negatively valenced images (Damasio et al., 1990). However, other studies have shown the opposite relationship: vmPFC activity has been found to negatively covary with skin conductance levels in healthy volunteers (Nagai et al., 2004), and vmPFC activity was negatively correlated with heart rate during a public speaking preparation task (Wager et al., 2009a; Wager et al., 2009b). These discrepancies could be due to the presence of different functional subregions of the vmPFC (Myers-Schulz and Koenigs, 2012), or could possibly reflect different qualities in a task such as attentional demands (Simpson et al., 2001).

Activity in the vmPFC has been considered important in top-down regulation of amygdala activity and anxiety, mainly owing to results of studies utilising explicit emotional regulation tasks (Etkin et al., 2011; Ochsner and Gross, 2005; Urry et al., 2006; Delgado et al., 2008). However, when no explicit instructions to regulate emotions are given, there is evidence to suggest that anterior vmPFC and amygdala co-activation occurs during processing of potential threat. For example, in a study in which healthy volunteers were tasked to ‘escape’ from a virtual predator, potential threat was associated with activity in the vmPFC and right basolateral amygdala, while during acute threat activity shifted to a midbrain network including the central amygdala and the periacqueductal grey matter (Mobbs et al., 2007). Further, adults with psychopathy exhibit reduced fear-potentiated startle in response to environmental threats, reduced amygdala responses to aversive emotional stimuli and reduced functional connectivity between the vmPFC and the amygdala (Blair, 2013). It is possible that increased connectivity between the vmPFC and the amygdala relates to a lower threshold to appraise aversive stimuli as threatening. This could explain why participants who show this pattern of activity when viewing aversive stimuli respond more convincingly to CO_2_ challenge.

GAD-7 scores were negatively associated at a trend level with functional connectivity between the MCC and the left amygdala when viewing angry compared with happy faces. A previous study has shown a similar relationship between MCC/amygdala connectivity and anxiety. In a sample of healthy adult men, functional connectivity between the left MCC and the left amygdala when viewing negative compared with neutral faces was negatively correlated with trait anxiety (Kienast et al., 2008). The MCC appears to play a complex role in selecting appropriate actions based on incoming negatively valenced information (Shackman et al., 2011; Vogt, 2014). This is supported by studies that show the MCC is activated by cognitive reappraisal of negative emotional stimuli (Ochsner et al., 2002), and when ‘escaping’ from a virtual predator (Mobbs et al., 2009; Mobbs et al., 2007). It is therefore possible that reduced functional connectivity between this region and the amygdala suggests reduced ability to engage appropriate control mechanisms to manage aversive emotional states.

Other regions important in the cognitive control of emotions include the lateral prefrontal cortices (Ochsner and Gross, 2005). The fMRI correlates of response to CO_2_ challenge have been explored in one previous study, which suggested that dorsolateral PFC activity was inversely related to the experience of anxiety during CO_2_ challenge and during unpredictable threat (Balderston et al., 2017). In this study, the authors utilised a no shock, predictable shock and unpredictable threat of shock (NPU) paradigm to identify anxiety-related brain activity in healthy volunteers. During a separate testing session, these participants also underwent CO_2_ challenge. Principle component and general linear model analyses showed a significant inverse relationship between panic-related symptoms such as heart rate and dizziness during CO_2_ challenge and anxiety-related dorsolateral PFC activity during unpredictable threat (Balderston et al., 2017). The dorsolateral PFC was not included as a region of interest in the current pilot study, as there were no significant activations in this region in our hypothesis-generating cohort. We need to explore in further studies whether activity in the dorsolateral PFC is functionally connected to the regions implicated in the current study.

Taken together, our findings suggest there are inter-individual differences in the function of negative affective valence networks and these relate to response to CO_2_ challenge. Increased connectivity between vmPFC and right amygdala, and reduced connectivity between MCC and left amygdala, when viewing aversive stimuli was associated with a greater anxiogenic response. A hypothesis which could explain these findings is that this pattern of activity occurs in participants with a lower threshold to assess aversive stimuli as threatening, and who have reduced ability to then manage this emotional response appropriately. We cannot test this hypothesis in the current dataset. However, further studies should be undertaken to test this hypothesis and to assess whether this translates into clinical populations, especially as trait markers of anxiety and degree of response to CO_2_ challenge might predict the development of anxiety symptoms (Fluharty et al., 2016; Olatunji et al., 2009; Schmidt and Zvolensky, 2007).

Our methods also had some limitations that need to be discussed. First, this was an exploratory pilot study with a small sample size and a relatively short behavioural task during the scan. For these reasons, we did not have the statistical power to carry out seed to voxel or whole-brain voxel to voxel PPI analyses. Second, we did not take measures of subjective mood or anxiety during the scanning session. This means we do not know whether participants experienced similar emotional states during scanning and during CO_2_ challenge. However, given that the network of regions activated by the faces task was consistent with the literature and included the amygdala, this suggests the task did indeed recruit negative valence systems: but our results need to be interpreted with caution due to these limitations.

## Conclusion

The purpose of this study was to explore the association between the functional architecture of negative affective valence networks and response to CO_2_ challenge in healthy volunteers. We found that increased functional connectivity between vmPFC and right amygdala, and reduced connectivity between MCC and left amygdala, when viewing angry faces was associated with anxiety during CO_2_ challenge. It is possible that this pattern is consistent with increased threat responsivity to aversive stimuli and a reduced capacity to appropriately manage the related emotional response. Further studies are required to test this hypothesis and to understand whether this translates into clinical populations. Functional connectivity within emotional processing networks could represent a potential biomarker for stratifying healthy volunteers and examining correlates of response in trials utilising experimental medicine models.

## Acknowledgements

We would like to thank Chris Everitt and Chris Watson, of University Hospital Southampton, Southampton, UK, and David Keep, of the Academic Unit of Psychology, University of Southampton, for all their help with data collection.

## Author contributions

Conceived and designed the experiments: NTMH, MJB, AD, MG, DSB. Performed the experiments: NTMH. Analysed the data: NTMH, MJB. Contributed reagents/materials/analysis tools: MJB, AD. Wrote the manuscript: NTMH, MJB, AD, MG, DSB.

